# Confusion will be my epitaph: Genome-scale discordance stifles phylogenetic resolution of Holothuroidea

**DOI:** 10.1101/2022.12.11.519962

**Authors:** Nicolás Mongiardino Koch, Ekin Tilic, Allison K. Miller, Josefin Stiller, Greg W. Rouse

## Abstract

Sea cucumbers (Holothuroidea) are a diverse clade of echinoderms found from intertidal waters to the bottom of the deepest trenches. Their reduced skeletons and limited number of phylogenetically-informative traits have long obfuscated morphological classifications. Sanger-sequenced molecular datasets have also failed to constrain the position of major lineages. Noteworthy, topological uncertainty has hindered a resolution for Neoholothuriida, a highly diverse clade of Permo-Triassic age. We perform the first phylogenomic analysis of Holothuroidea, combining existing datasets with twelve novel transcriptomes. Using a highly-curated dataset of 1,100 orthologues, our efforts recapitulate previous results, struggling to resolve interrelationships among neoholothuriid clades. Three approaches to phylogenetic reconstruction (concatenation under both site-homogeneous and site-heterogeneous models, and coalescent-aware inference) result in alternative resolutions, all of which are recovered with strong support, and across a range of datasets filtered for phylogenetic usefulness. We explore this intriguing result using gene-wise log-likelihood scores, and attempt to correlate these with a large set of gene properties. While presenting novel ways of exploring and visualizing support for alternative trees, we are unable to discover significant predictors of topological preference, and our efforts fail to favor one topology. Neoholothuriid genomes seem to retain an amalgam of signals derived from multiple phylogenetic histories.

## Introduction

Holothuroidea, commonly known as sea cucumbers, is arguably the most morphologically diverse major clade of extant Echinodermata (Fig. 1). The smallest adults can be less than 1 cm in length, as seen in the meiofaunal *Leptosynapta minuta* [1] and epibenthic *Incubocnus* [2]. The largest can be thin and elongate, reaching several meters length, as in the snake sea cucumber *Synapta maculata* [3], or they may be less than a meter but robust and weighing over 5 kg, as in the case of *Holothuria fuscopunctata* [4]. While predominantly benthic as adults, some taxa are capable of swimming and there are even forms that spend their entire lives in the water column [5]. While all holothuroids have a ring of tentacles and are deposit or filter feeders, some clades lack tube feet and have a substantially reduced water vascular system, traits otherwise developed across all echinoderms. They can also entirely lack calcareous elements (ossicles) in the body wall, or these can be expanded to form overlapping plates that build a rigid test [6]. There are currently 1,775 accepted extant species of holothuroids [7] found in ocean waters that range from the intertidal to the bottom of the deepest trenches [8, 9]. Especially in benthic deep-sea habitats, they can constitute the vast majority of total biomass and have a strong impact on ecosystem functioning, bioturbation, and nutrient cycling [10-12].

**Figure 1:**
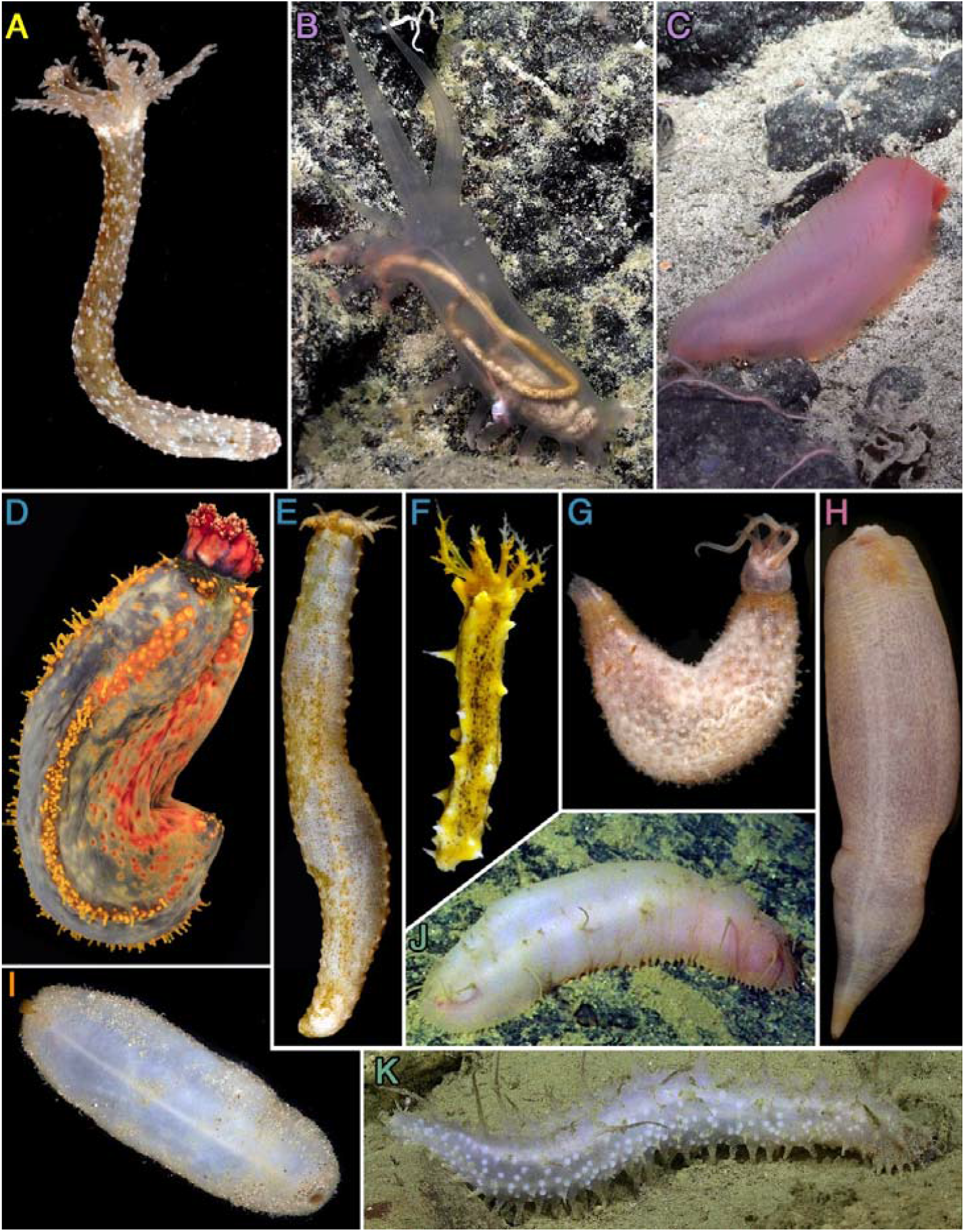
Representative holothuroid diversity included in this study. **A**. *Synapta* sp. **B**. *Peniagone* cf. *vitrea*. **C**. *Benthogone* sp. **D**. *Pseudocolochirus violaceus*. **E**. *Abyssocucumis albatrossi. F. Colochirus robustus*. **G**. *Ypsilothuria* n. sp. (SIO-BIC E6221). **H**. *Molpadia amorpha*. **I**. *Pseudostichopus* sp. **J**. Synallactidae. **K**. *Bathyplotes* cf. *moseleyi*. The classification of these terminals can be found in Table 1; colors for panel letters follow those used for major lineages in Figure 2. All photos except **G** are of the voucher specimens sequenced (catalogue numbers can be found in Table 1; further sampling information is accessible through the SIO-BIC online database, https://sioapps.ucsd.edu/collections/bi/).

While multiple morphological attempts have been made to delineate major subdivisions within Holothuroidea, these have been limited by the extreme simplification of their skeleton (relative to other echinoderms), the delicate and fragile nature of their bodies, and the small number of traits that provide useful information at high taxonomic levels [13-15]. The most recent revision of the group’s classification was based on a six-gene dataset including terminals from 25 of the 29 accepted family-ranked taxa [16]. This study recovered a basal split within sea cucumbers between Apodida, a clade characterized by a complete loss of tube feet, and Actinopoda (among which secondary reductions or loss of tube feet occur only within Molpadida). Actinopoda were further subdivided into Elasipodida and Pneumonophora, the latter of which includes all species with respiratory trees, a unique cloacal invagination that plays an important (although not exclusive) role in respiration [17, 18]. Furthermore, the names Holothuriida and Neoholothuriida were applied to the main subdivisions within Pneumonophora, with four well-supported major clades inside Neoholothuriida: Dendrochirotida, Molpadida, Persiculida, and Synallactida. However, the relationships among these four lineages remained uncertain. A phylogenetic resolution for the major neoholothuriid lineages is necessary to explore the origins of the high morphological and ecological disparity harbored by this clade, as well as to establish a natural classification framework for a substantial fraction of sea cucumber diversity (62% of species-level diversity is contained within Neoholothuriida [7]). Miller et al. [16] concluded that meeting these objectives would likely require sequencing efforts of a different magnitude.

**Table 1:**
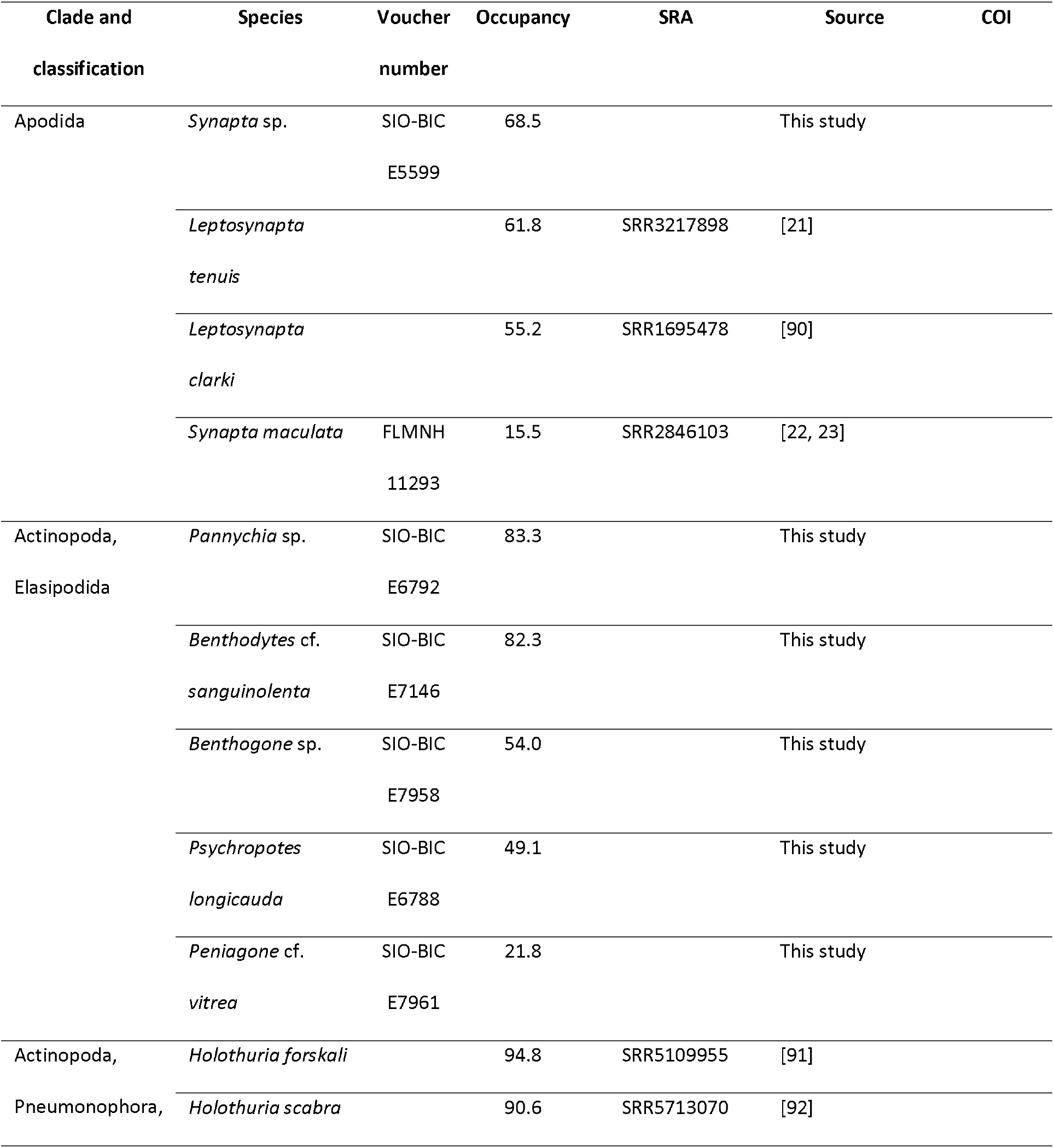
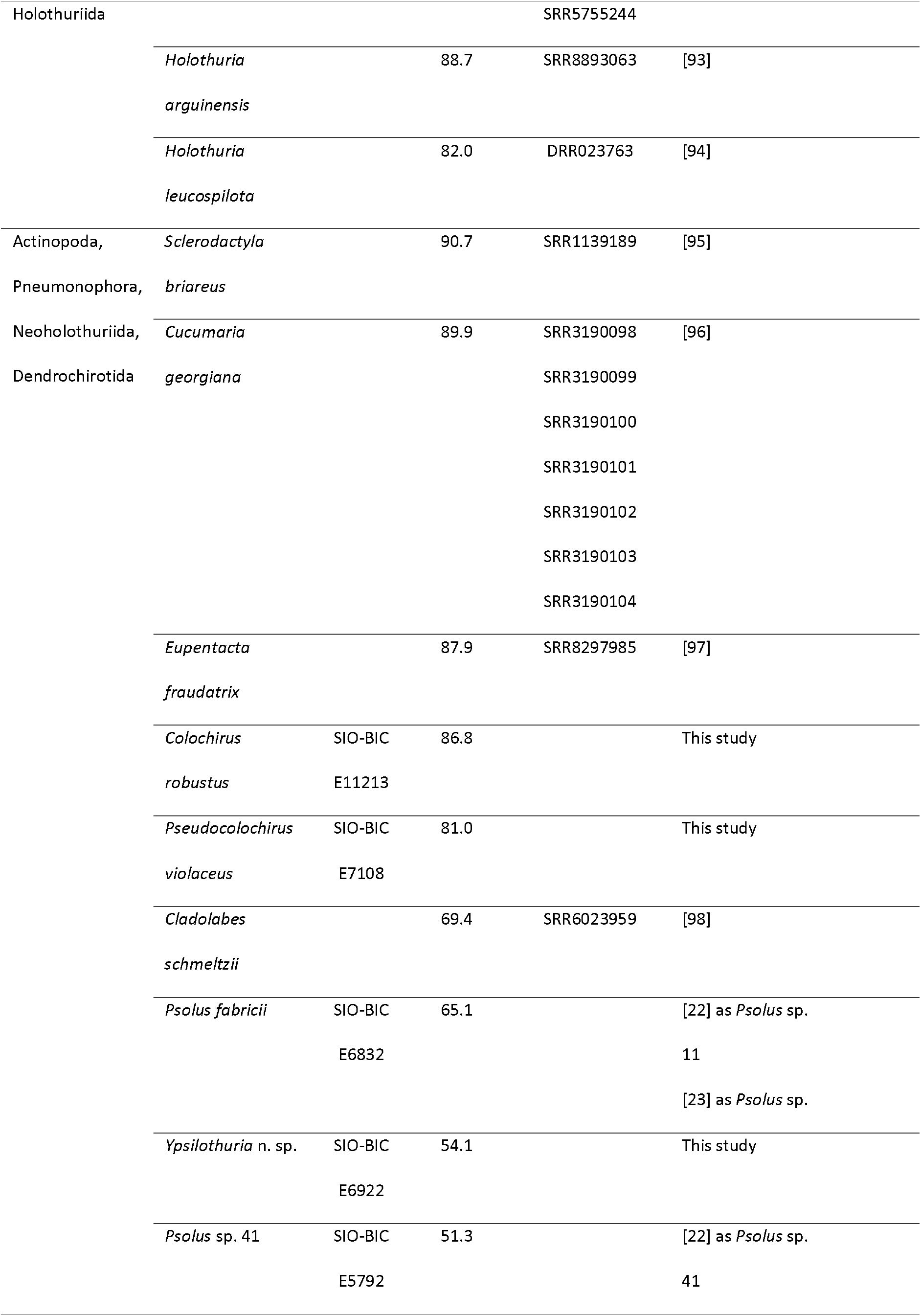
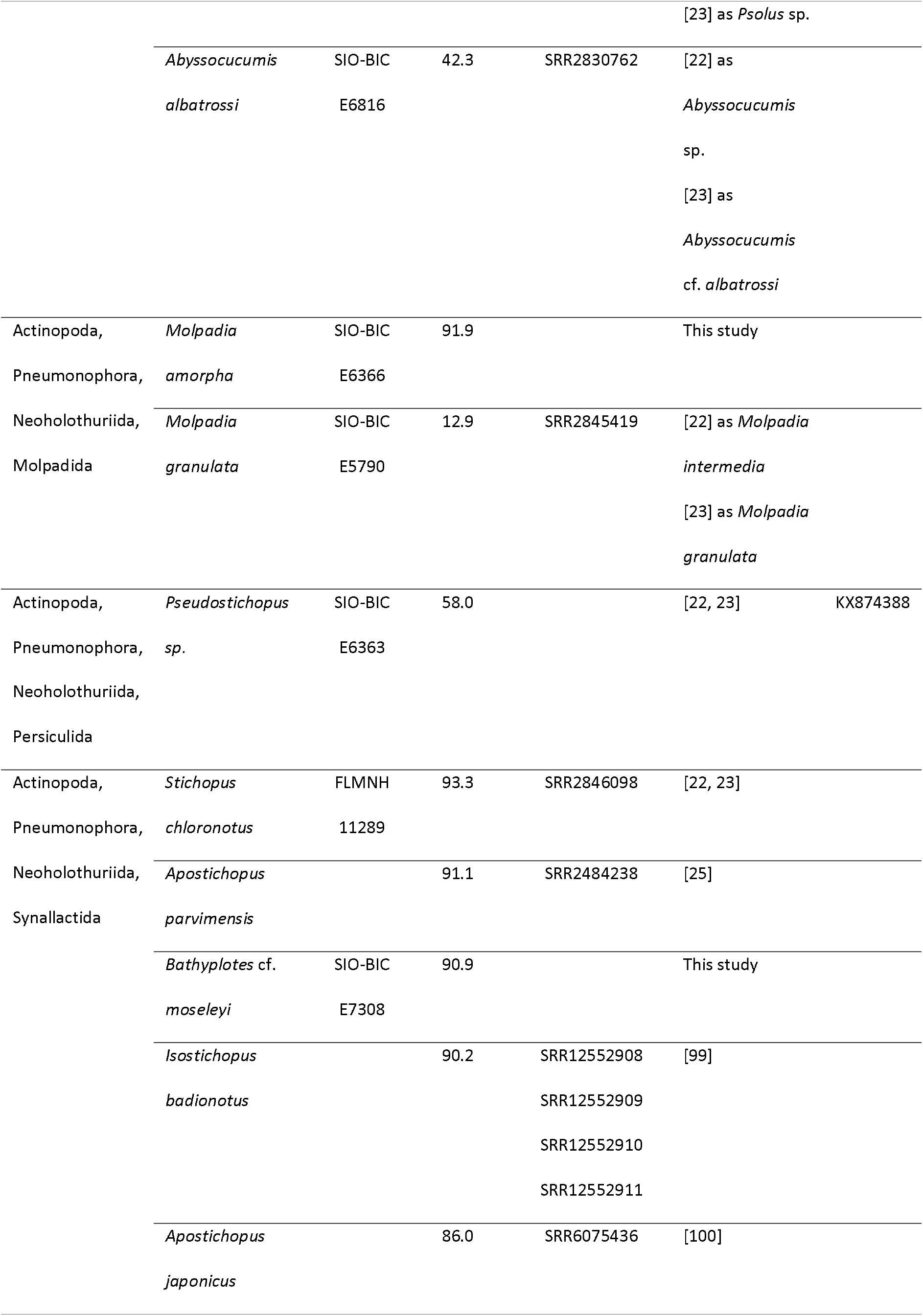
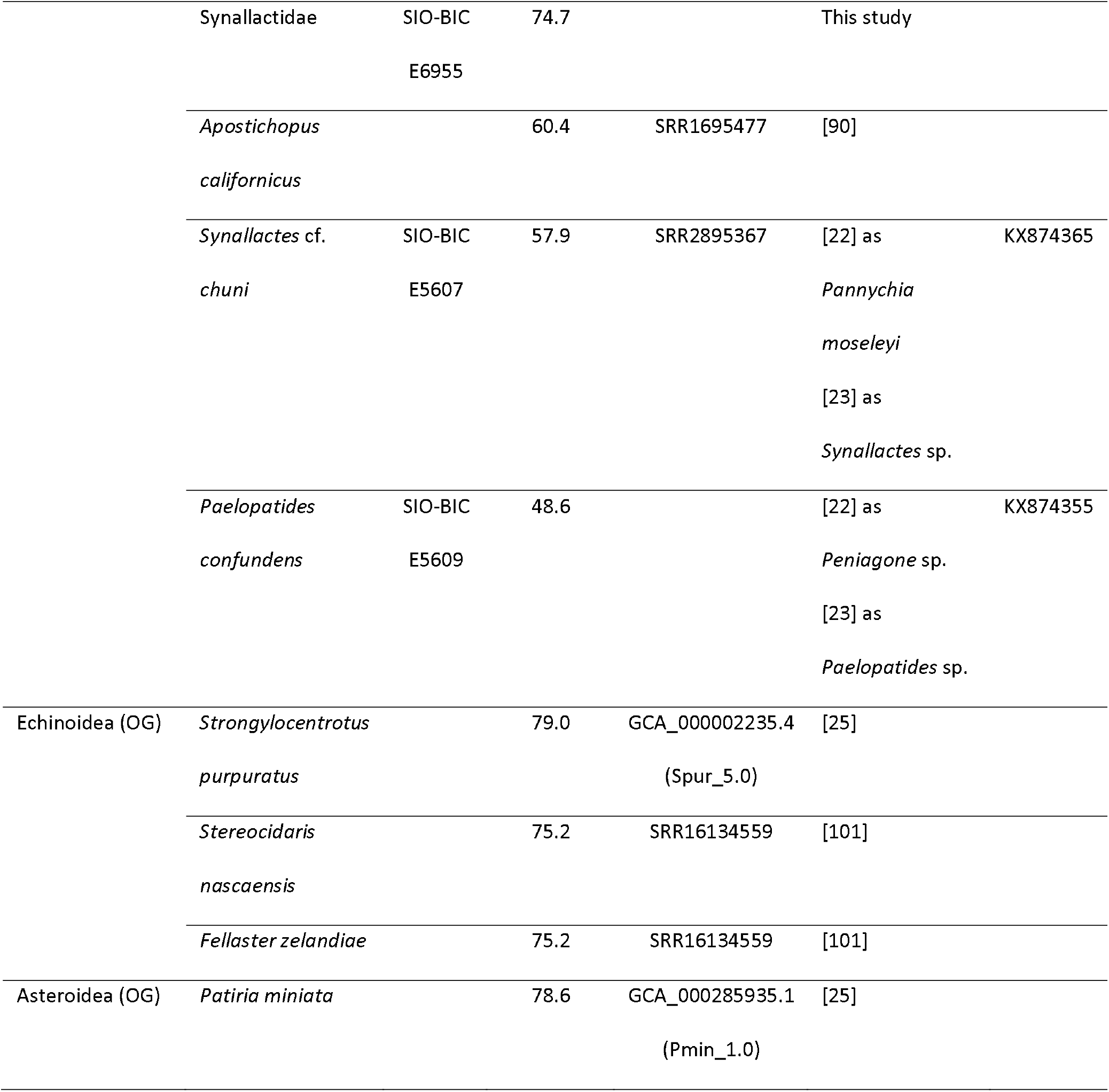
Terminals included in the phylogenomic dataset. Voucher numbers are provided for all novel transcriptomic datasets, as well as for those available on EchinoDB [23]. For the latter, some changes in species identifications are noted. NCBI accession numbers of COI sequences are noted when available. Higher-level classification follows Miller et al. [16]. Within major clades, terminals are ordered by their occupancy level. OG = outgroups.

Here we present the first phylogenomic study of sea cucumbers, the last major eleutherozoan clade (which further includes echinoids [19], asteroids [20], and ophiuroids [21]) to have its phylogeny tackled using genome-scale datasets. Through the generation of novel transcriptomic resources for holothuroids we built a molecular dataset encompassing over a thousand orthologs. The goal was to resolve some of the lingering uncertainties in the holothuroid tree of life, yet a continuing lack of resolution encouraged novel ways to explore phylogenomic datasets.

## Materials and methods

### Taxon sampling, extraction and sequencing

Sea cucumber specimens were collected by SCUBA diving, snorkeling, dredging, and remotely operated vehicles (ROV), or purchased from aquarium suppliers, and vouchers were deposited at the Benthic Invertebrate Collection, Scripps Institution of Oceanography (SIO-BIC, see Table 1). Species identification was based on multiple lines of evidence, including anatomical (gross and ossicle morphology), biogeographical and molecular (mitochondrial cytochrome c oxidase subunit I, COI) information. DNA extractions and COI amplifications followed protocols described in Miller et al. [16], and sequences are deposited in NCBI (accession numbers available in Table 1). Previous identifications of transcriptomic vouchers at SIO were also revised, including those sequenced and released as part of EchinoDB [22, 23].

For large specimens, tissue was dissected from the body wall or tube feet, while whole body sections were sampled for the remainder. Sampled tissues were finely chopped, placed in RNA*later* (Invitrogen) buffer solution, and stored at -80 °C. RNA extractions were performed from Trizol (Thermofisher), using Direct-zol RNA Miniprep Kit with in-column DNAse treatment (Zymo Research). mRNA was isolated with Dynabeads mRNA Direct Micro Kit (Invitrogen). mRNA concentration was estimated using Qubit RNA broad range assay kit (Thermofisher), and quality was assessed using RNA ScreenTape with an Agilent 4200 TapeStation on an Agilent Bioanalyzer 2100. Most libraries were prepared using a KAPA-Stranded RNA-Seq kit targeting a 200-300 bp insert size, and results were assessed using DNA ScreenTape (Bioanalyzer 2100). Libraries were then sequenced in multiplexed (8 libraries per lane) pair-end runs using 150 bp paired end Illumina HiSeq 4000 at the UC San Diego IGM Genomics Center. To minimize read crossover, we employed 10 bp sequence tags designed to be robust to indel and substitution errors [24]. For four samples (*Benthodytes* cf. *sanguinolenta, Benthogone* sp., *Colochirus robustus*, and *Peniagone* cf. *vitrea*), library preparation and multiplexed pair-end sequencing on an Illumina NovaSeq 6000 PE150 was performed by Novogene.

Twelve novel transcriptomes were generated for this study and combined with publicly available genomic and transcriptomic datasets downloaded from NCBI and EchinoBase [25]. Final taxonomic sampling included 35 holothuroids as well as three echinoids and one asteroid outgroups (Table 1). Raw files for all novel datasets, as well as those so far available only on EchinoDB [23], are deposited in the NCBI sequence read archive (SRA) under Bioproject XXX. All assemblies are available at the Dryad data repository YYY.

### Assembly, sanitation and matrix construction

Reads were trimmed or excluded based on quality scores using Trimmomatic v 0.3.6 under default settings [26]. Additional sanitation steps were implemented by the Agalma 2.0 pipeline [27], resulting in the removal of reads based on compositional and quality filtering criteria, as well as those mapping to rRNA sequences or retaining adapter sequences. Remaining reads were assembled *de novo* with Trinity v. 2.5.1 [28]. Assemblies were then screened for contaminants using alien_index v 3.0 [29]. Transcripts with substantially better BLAST+ [30] hits to a dataset of well-curated archaeal, bacterial and fungal genomes than to a metazoan database (both available from http://ryanlab.whitney.ufl.edu/downloads/alien_index/), defined as those exhibiting an alien index > 45 (see [31]), were excluded. Sanitized transcriptomes were imported back into Agalma for orthology inference [27, 32], alignment using MAFFT v. 7.305 [33], and quality-based trimming with GBLOCKS v. 0.91b [34]. The resulting supermatrix was reduced using a 70% occupancy threshold, resulting in a dataset composed of 1,159 orthologs coded as amino acids (from a total of 13,767).

Gene trees were inferred from each amino acid alignment with ParGenes v. 1.0.1 [35], using optimal models and 100 bootstrap replicates. These were analyzed with TreeShrink v. 1.3.1 [36] (using parameters `-q 0.01 -k 3 -b 25`), which employs taxon-specific distributions of root-to-tip distances to identify outlier sequences potentially suffering from errors in alignment or orthology inference. Identified outliers were removed from both gene trees and individual alignments, and a new supermatrix was concatenated. As a final sanitation step, the data was run using *genesortR* [37, 38], which ordered all loci based on decreasing estimates of phylogenetic usefulness (Fig. S1). The worst-ranked 59 loci (5.1% of supermatrix) were further discarded, resulting in a final dataset of 1,100 loci and 264,991 amino acid positions. Two smaller datasets, composed of the top-scoring (i.e., most phylogenetically useful) half and quarter of loci (550 and 225, respectively) were also output for analysis.

### Phylogenetic inference and signal dissection

Phylogenies were inferred from all three datasets using a variety of approaches. First, gene trees were provided to the coalescent-aware summary method ASTRAL-III [39], which employs local posterior probabilities to estimate node support [40]. Second, tree inference was performed under a best-fit partitioned model with IQ-TREE 2 v. 2.1.3 [41-43], using the fast-relaxed clustering algorithm to merge individual loci (using parameters `-m MFP + MERGE -rclusterf 10 -rcluster-max 3000`). Finally, the site-heterogeneous model GTR+CAT-PMSF [44] was used as an efficient alternative to the computationally-demanding CAT model family [45]. For each dataset, short runs of 1,100 generations were done in PhyloBayes-MPI v. 1.8.1 [46] under a fixed topology (that obtained with ASTRAL-III) to approximate site-specific stationary distributions and amino acid exchangeabilities under the GTR+CAT model. Model parameters were summarized after discarding the initial 100 generations as burn-in, and reformatted using scripts available at https://github.com/drenal/cat-pmsf-paper. Tree inference was then performed in IQ-TREE 2 under maximum likelihood using the PMSF method [47], setting exchangeabilities and site-specific frequencies to the posterior mean estimates previously obtained with PhyloBayes. For both concatenation approaches, support was estimated using 1000 replicates of ultrafast bootstrap [48].

Given persistent discordance among methods of inference regarding the resolution of major neoholothuriid clades, phylogenetic signal for the alternative topologies was explored using site-wise log-likelihood scores. Scores were computed with IQ-TREE 2 under the same best-fit partitioned model mentioned above, and using three constrained topologies differing only in the position of Molpadida, with the remaining incongruency fixed to the preferred resolution (see Results). Gene-wise log-likelihood scores were obtained through addition across alignment positions, and their differences for all pairs of topologies—known as ΔGLS values—were computed. Given linear dependency between the three ΔGLS values (∆*GLSx* = ∆*GLSy* - ∆*GLSz*), these were visualized in a two-dimensional space which was rotated using principal components analysis (PCA). A nominal ΔGLS threshold of ± 2 log-likelihood units was used to categorize loci as either informative or uninformative with regards to a given topological comparison.

To explore the drivers of differences in ΔGLS across loci, fifteen gene properties were estimated and treated as potential determinants. These were all calculated by *genesortR* [38], and included commonly used metrics of phylogenetic signal, potential sources of systematic bias, and estimates of the overall information content and evolutionary rate of each individual loci. Further details on these metrics can be found in Table S1.

Potential links between gene properties and the phylogenetic support for alternative neoholothuriid relationships were explored using two different statistical approaches. Associations between the PCA axes derived from ΔGLS values (representing major aspects of phylogenetic signal for competing hypotheses) and explanatory variables were first tested using the ‘envfit’ function in R package *vegan* [49]. This approach overlayed vectors onto the ordination plot depicting the directions and magnitudes of maximum correlation between individual gene properties and PCA scores. Each predictor was analyzed separately, and the significance of the correlations tested using 10,000 random permutations. Given the possibility of non-linear relationships between predictor and response variables, a second approach was explored in which ΔGLS values were transformed into a single categorical factor with three levels. For this, loci were categorized into: A) uninformative, including those for which all ΔGLS were within ±2 log-likelihood units; as well as those either B) supporting, or C) rejecting the resolution obtained using ASTRAL-III, defined as exhibiting at least one comparison favoring or disfavoring such topology, respectively, by an absolute ΔGLS value > 2. This categorization is supported by analyses showing that the main aspect of differences in phylogenetic signal across loci relates to their support for/against the topology obtained with ASTRAL-III, with very little ability to discriminate between the other alternatives (see Results). A conditional inference classification tree was fit to the data using function ‘ctree’ in R package *partykit* [50], assessing whether partitioning the data by values of any of the gene properties was able to generate subsets of loci that show similar topological preferences. A Bonferroni correction for multiple comparisons was applied, and significant predictors were visualized on the ordination plot using smooth surfaces fit using penalized regression splines [51].

All statistical analyses were performed in the R environment [52] using code reliant on the packages mentioned above, as well as *adephylo* [53], *ape* [54], *phangorn* [55], *phytools* [56], and those included in the *tidyverse* [57].

## Results

Phylogenetic inference under all methods explored and for the three datasets of different sizes recovered highly congruent and well-supported topologies (Figs. S2-4), which were also in broad agreement with the most recent large-scale study based on Sanger-sequenced loci [16]. As summarized in figure 2A, Apodida, Elasipodida and Holothuriida formed successive and monophyletic sister groups to the remainder of sea cucumber diversity included within Neoholothuriida. The latter was further subdivided into four major lineages: Dendrochirotida, Molpadida, Persiculida and Synallactida. Nodes defining all aforementioned clades had maximum support across analyses.

**Figure 2:**
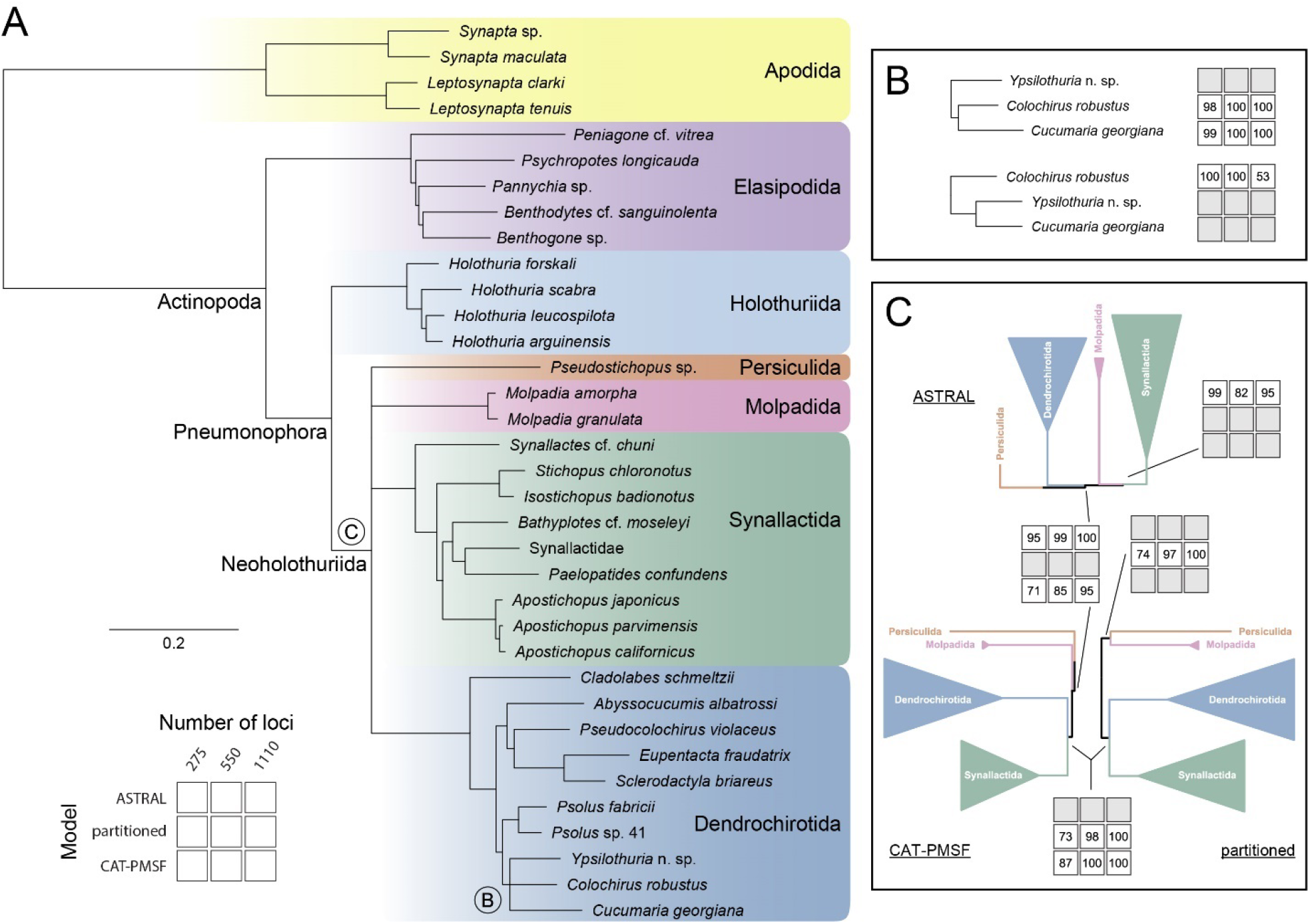
Summary of phylogenetic inference results. **A**. Strict consensus of the nine inference conditions explored, varying both the number of loci and the method of inference. Nodes disagreeing between analyses are collapsed and labeled (see panels **B-C** for further details); branch lengths are otherwise taken from the CAT-PMSF analysis of the full dataset. **B**. Monophyly of a clade composed of two cucumariid terminals, *Colochirus robustus* and *Cucumaria georgiana*, is rejected by ASTRAL-III, but upheld by the other methods (legend for support value grid is shown in **A**). **C**. Systematic disagreement between all methods of inference regarding relationships among major neoholothuriid clades. The resolution favored by each method is found across datasets of different sizes. Topologies, branch lengths and support values for each individual analysis are shown in Figs. S2-4.

Only two regions of the tree topology showed incongruent resolutions among the analyses performed (Figs. 2B-C). First, ASTRAL-III rejected a close relationship among two of the cucumariid species sampled, *Colochirus robustus* and *Cucumaria georgiana*, which otherwise formed a clade under concatenation approaches (Fig. 2B). Given the otherwise unambiguous support for a close relationship between *Colochirus* and *Cucumaria*, as well as the poor node support for the ASTRAL-III topology when using the complete dataset, we tentatively favor here the results obtained under concatenation methods. We note, however, that a monophyletic Cucumariidae was not recovered by our analyses regardless of how these terminals are resolved, as they were only distantly related to the remaining cucumariids (*Abyssocucumis, Pseudocolochirus*). In fact, discrepancies between our trees and the current family-level classification of sea cucumbers are pervasive, and also included the non-monophyly of elasipodid families Psychropotidae (*Psychropotes, Benthodytes*) and Laetmogonidae (*Benthogone, Pannychia*), the dendrochirotid family Sclerodactylidae (*Cladolabes, Eupentacta, Sclerodactyla*), and the synallactid families Synallactidae (*Synallactes, Bathyplotes, Paelopatides*) and Stichopodidae (*Stichopus, Isostichopus, Apostichopus*).

A second and more striking topological discordance involved the organization of the four major lineages within Neoholothuriida (Fig. 2C). Each one of the different methods of inference proposed an alternative resolution for the clade, which were recovered regardless of dataset size and strongly supported (values > 95) when employing the complete supermatrix. While all three inference methods agreed on a subtree in which Dendrochirotida and Synallactida share a closer relationship than either one does with Persiculida, the position of Molpadida within this scaffold was highly unstable and methodologically sensitive. Supported alternatives included a placement of Molpadida as sister to either Synallactida, Persiculida, or Synallactida + Dendrochirotida (henceforth referred to as ‘ASTRAL’, ‘partitioned’, and ‘CAT-PMSF’ topologies, respectively). Despite this level of uncertainty, our analyses still reject the long-hypothesized close relationships between Molpadida and Dendrochirotida [13, 15], as well as the topology of Miller et al. [16] in which Dendrochirotida placed as sister group to all other neoholothuriids.

To further explore the phylogenetic signal for competing neoholothuriid topologies, we estimated gene-wise log-likelihood scores for the three alternative resolutions of this clade. A PCA of differences in the scores obtained for pairs of trees (ΔGLS) revealed that the topological preferences of loci could be summarized using a single major underlying axis which accounted for 85.3% of total variance (Fig. 3A). The scores of loci along this first PC axis represented the relative support either for or against the ASTRAL topology (Fig. 3B). On the other hand, the ability of loci to discern between the partitioned and CAT-PMSF trees was much weaker and mainly captured by the second PC axis, which explained only 14.7% of variance. The absolute values of ΔGLS were generally small, with most loci (615 loci, 55.9% of the complete dataset) being relatively uninformative regarding relationships among neoholothuriid clades (Figs. 4A and S5). Nonetheless, the remainder of the dataset was once again roughly evenly split into a fraction that supported the ASTRAL configuration (207 loci, 18.8%), and one that rejected it in favor of either one, or both, of the topological alternatives (278 loci, 25.3%).

**Figure 3:**
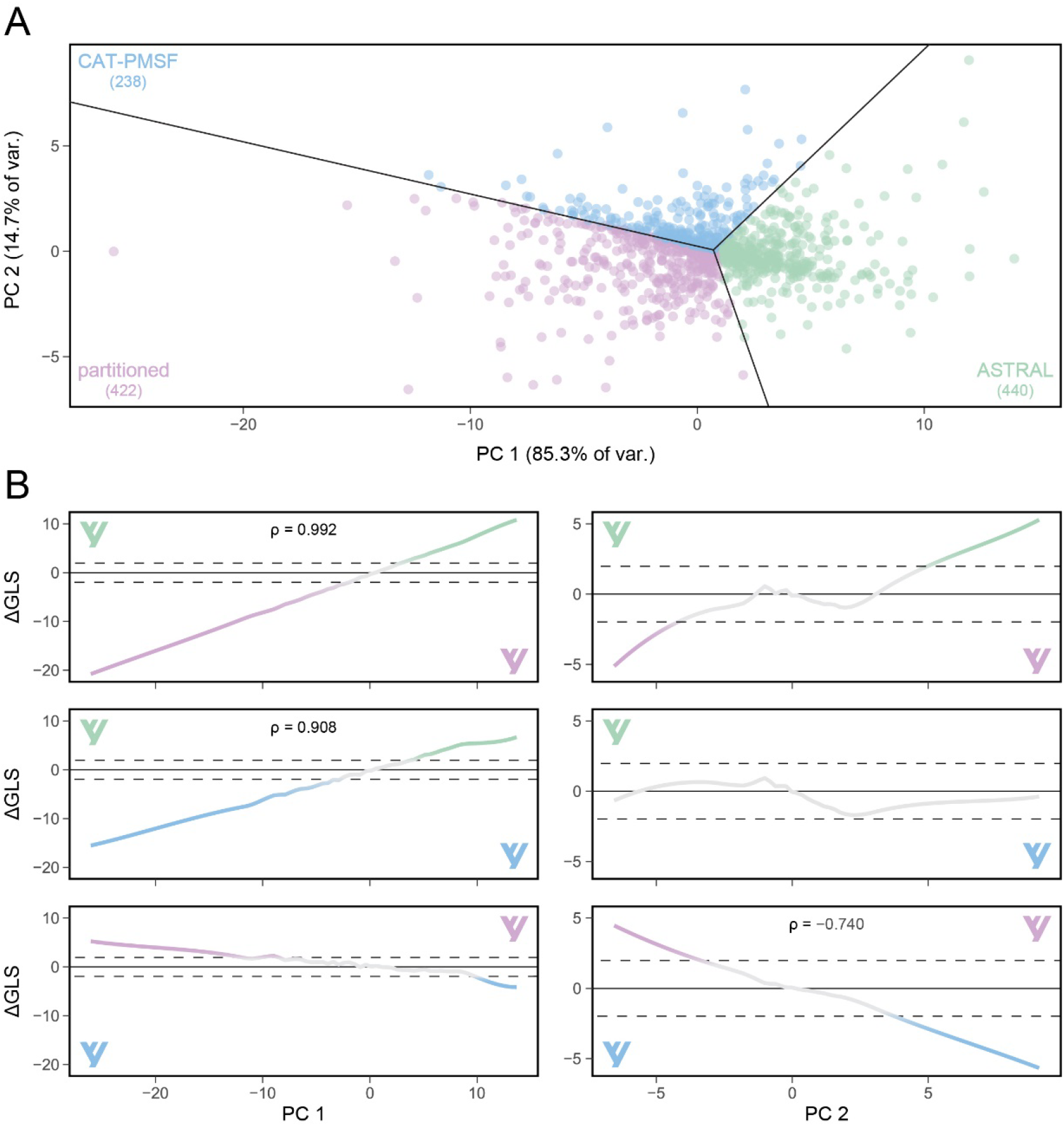
Exploration of support for alternative neoholothuriid topologies across loci. **A**. Principal components (PC) axes obtained from the three ΔGLS. Percentages of explained variance are shown on axis labels. Loci are color coded depending on their favored topology. **B**. Relationship between the PC axes and the scores of individual ΔGLS. Trendlines correspond to LOESS smoothing curves, and ρ values (Spearman’s rank correlation coefficients) are shown when absolute values > 0.7, taken to represent strong correlations. The area included within ± 2 log-likelihood units is highlighted, and considered an area of weak support. Note the markedly different scales of the y-axes for PCs 1 and 2. Topologies are color coded as in A.

**Figure 4:**
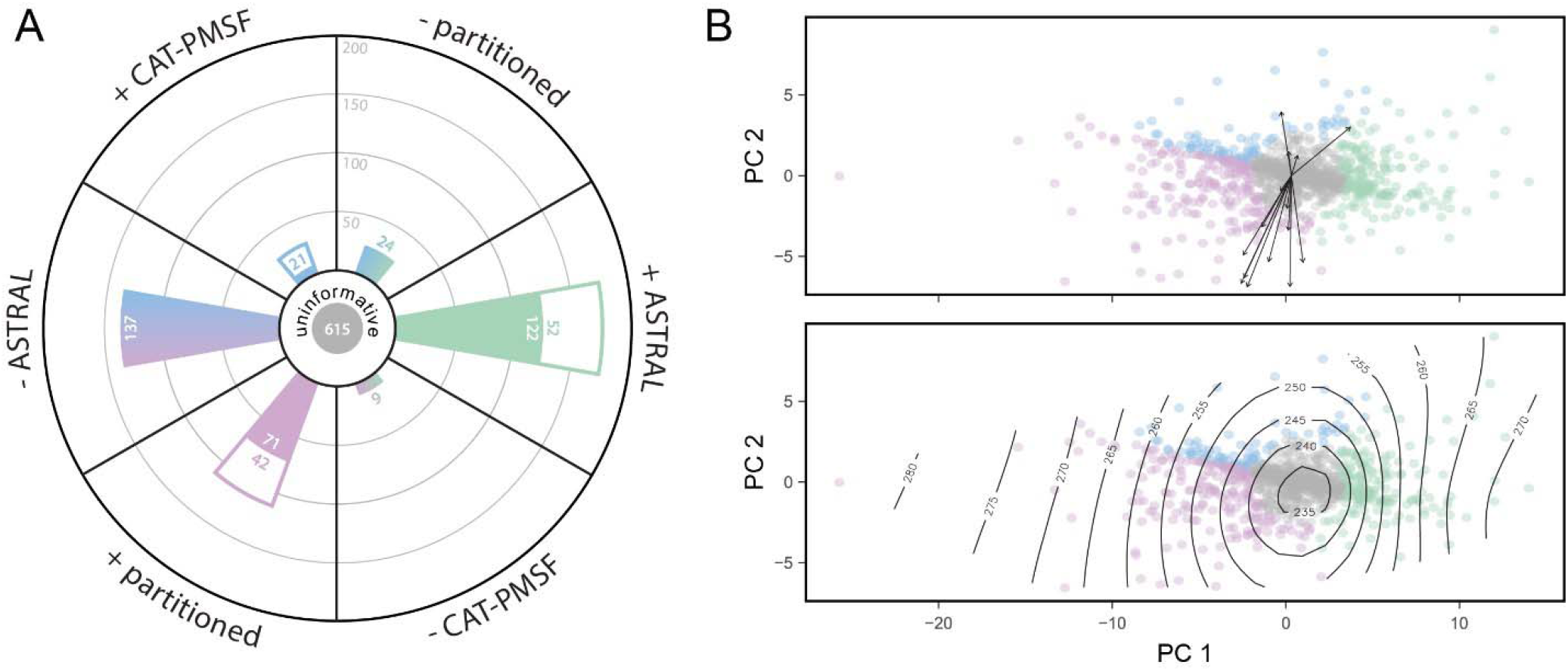
Categorization of loci depending on their favored topology, and exploration of potential determinants. Coloring scheme follows that of Figure 3. **A**. Most loci (615, 55.9% of the full dataset) can be considered uninformative regarding relationships among neoholothuriid clades. The remainder can be classified into those supporting a given topology (denoted using a +) if they favor a given resolution against both alternatives (colored section of bar chart) or only one (white section of bar chart) with a ΔGLS ≥ 2; or rejecting a given topology (denoted using a -). The number of loci either supporting (right side of wheel) or rejecting (left side of wheel) the ASTRAL topology are comparable in number: 207 (18.8%) vs. 278 (25.3%), respectively. Further details on loci categorization can be found in Figure S5. **B**. Top: Exploration of 15 potential determinants of ΔGLS. Arrows indicate directions of maximum correlation between scores and determinants; their length is scaled to the strength of the correlation. Predictors mostly load onto PC 2. R^2^ and *p*-values are shown in Table S1, but no correlation is significant. Bottom: Smoothed surface of alignment length, the only significant determinant found using a classification tree. Longer loci are more likely to be informative, yet alignment length does not predict which topology is preferred (see Fig. S6).

None of the fifteen gene properties explored was recovered as a significant predictor of ΔGLS (Fig. 4B). Furthermore, these metrics correlated mostly with PC 2 (Table S1), leaving the major aspect of topological preference entirely unexplained. An alternative approach based on classification trees recovered one significant predictor: uninformative loci were significantly more likely to have a short alignment length, but this property also fails to explain which resolution was preferred by longer and more informative loci (Fig. S6).

## Discussion

Phylogenetic incongruence is a hallmark of genome-scale datasets [58-60]. A wide range of biological processes and methodological artifacts can lead phylogenomic datasets to harbor a mixture of phylogenetic signals, which can be differentially amplified by methods of reconstruction to produce conflicting, yet well-supported, topologies [61-63]. Different avenues have been proposed to ameliorate phylogenetic incongruence and favor a specific resolution for recalcitrant nodes. One strategy is to focus on data filtering, exploring the effects of removing sites and/or loci with unexpectedly high topological preferences [64, 65], or those showing evidence of contributing mostly phylogenetic noise or biases [38, 66]. Alternatively, methods have been developed to dissect alternative signals [67-69] in the hopes that one emerges as a better-justified option. Finally, exploring a range of inference methods, which vary in their realism, complexity, susceptibility to errors, and (potentially) relative fit, can also be used to justify favoring one among several alternative hypotheses [70-72]. However, even after exhaustive testing of these options, a robust resolution for some nodes on the tree of life remains elusive [73-76], awaiting the discovery of novel phylogenetic markers, improved taxon sampling, or methodological developments.

We propose here that the early diversification of neoholothuriid sea cucumbers, an ancient, diverse, and morphologically heterogeneous clade, constitutes another example of a group that defies phylogenetic resolution. Previous studies had acknowledged that a robust topology for Neoholothuriida was likely unattainable with the use of small molecular datasets [16], yet phylogenetic resolution remains out of reach even when employing more than a thousand loci. The reason underlying this uncertainty is not a lack of statistical power, but the presence of multiple signals supporting alternative trees. While a node uniting Dendrochirotida and Synallactida to the exclusion of Persiculida emerges from all our analyses, the position of Molpadida within this topology remains uncertain. A coalescent-aware method of reconstruction places Molpadida inside the clade containing Dendrochirotida and Synallactida, while concatenation-based methods place it outside, with further disagreement emerging depending on whether site-homogeneous or site-heterogeneous models are used. All three of these topological alternatives are well-supported and robust to gene subsampling, and thus represent an example of remarkable methodological sensitivity. Further exploration reveals that our dataset is unlikely to contain enough information to disambiguate between the topologies supported by alternative concatenation methods. On the other hand, the placements of Molpadida either inside or outside of the node containing Dendrochirotida + Synallactida are each strongly supported by substantial fractions of the data (19 and 25% of loci, respectively).

Complex site-heterogeneous models, such as the CAT family, are likely to fit genome-scale datasets better [47, 77, 78], but the use of model fit statistics when comparing mixture models against other alternatives (such as partitioned models) has been criticized [79]. Furthermore, issues relating to convergence, missing data, and over-parameterization [80-82] have still led many to question the results obtained under CAT models. Similarly, coalescent-aware methods have outperformed concatenation in a number of simulation scenarios [83, 84], yet doubts remain regarding their usefulness to resolve ancient divergences, given that gene tree error is expected to surpass incomplete lineage sorting as the dominant source of incongruence for deep nodes [85]. The fit of summary methods (such as ASTRAL) is also impossible to evaluate relative to that of others, further complicating arriving at an objective way of preferring one method of inference from among those tested here.

In the absence of clear guidance as to which inference method can be preferred, we focused instead on evaluating the amount and quality of the signals supporting alternative placements of Molpadida. We used ΔGLS as proxies for the topological preference of loci (as in e.g., [19, 64, 86]), extending this type of analysis to simultaneously consider three alternative topologies. This allowed us to uncover a strong asymmetry in the ability of loci to distinguish between alternative trees and, as explained above, redirect our efforts to assessing two broad topological alternatives. Although many studies have succeeded in disentangling phylogenetic from non-phylogenetic signals by exploring loci quality [87-89], our attempts failed to find any determinants of topological preference: loci supporting alternative positions of Molpadida do not differ in their levels of phylogenetic signal, systematic biases, amounts of information or evolutionary rates. The only major pattern uncovered is that longer loci are more likely to harbor some sort of signal, a predictable and relatively trivial result stemming from the increased statistical power of longer alignments. Given a lack of evidence showing that competing hypotheses can be ascribed to systematic biases, we tentatively conclude that biological processes possibly underlie phylogenetic incongruence in this case, and that neoholothuriid evolution might be better explained by ancient events of reticulation, as produced by processes such as ancient hybridization and incomplete lineage sorting.

Although the exact placement of Molpadida remains challenging to ascertain, phylogenomics reveals an otherwise robust higher-level topology for sea cucumbers. These efforts are major step towards a stable classification for the group, corroborating much of the most recent classification based on molecular data [16]. The results presented here constitute a necessary tool with which to elucidate the times of origin, morphological evolution and diversification dynamics of one of the major lineages of marine invertebrates. At the same time, they also show the extent to which the current family-level classification scheme of holothuroids is at odds with their evolutionary history, highlighting the need for phylogenomic investigations with much-expanded taxon sampling and consequent morphological reassessments.

## Acknowledgements

Many thanks to Bob Vrijenhoek and David Clague of the Monterey Bay Aquarium Research Institute and the captain and crew of the R/V *Western Flyer* for sampling opportunities. Thanks also to the Schmidt Ocean Institute and chief scientists, Erik Cordes, Pete Girguis, and Lisa Levin and science parties of the R/V *Falkor* cruises FK181005, FK190106, and FK210726. We are also grateful to the captain and crew of the R/V *Robert Gordon Sproul* and the R/V IB *Nathaniel B. Palmer* and science participants during the cruise NBP-1303. Many thanks to the pilots of the remote operated vehicles *Doc Ricketts* and *SuBastian* for crucial assistance in specimen collection. We also thank Avery Hiley and Marina McCowin for assistance in the laboratory and to Charlotte Seid (SIO-BIC) for specimen cataloguing.

## Supplementary Information

**Figure S1:**
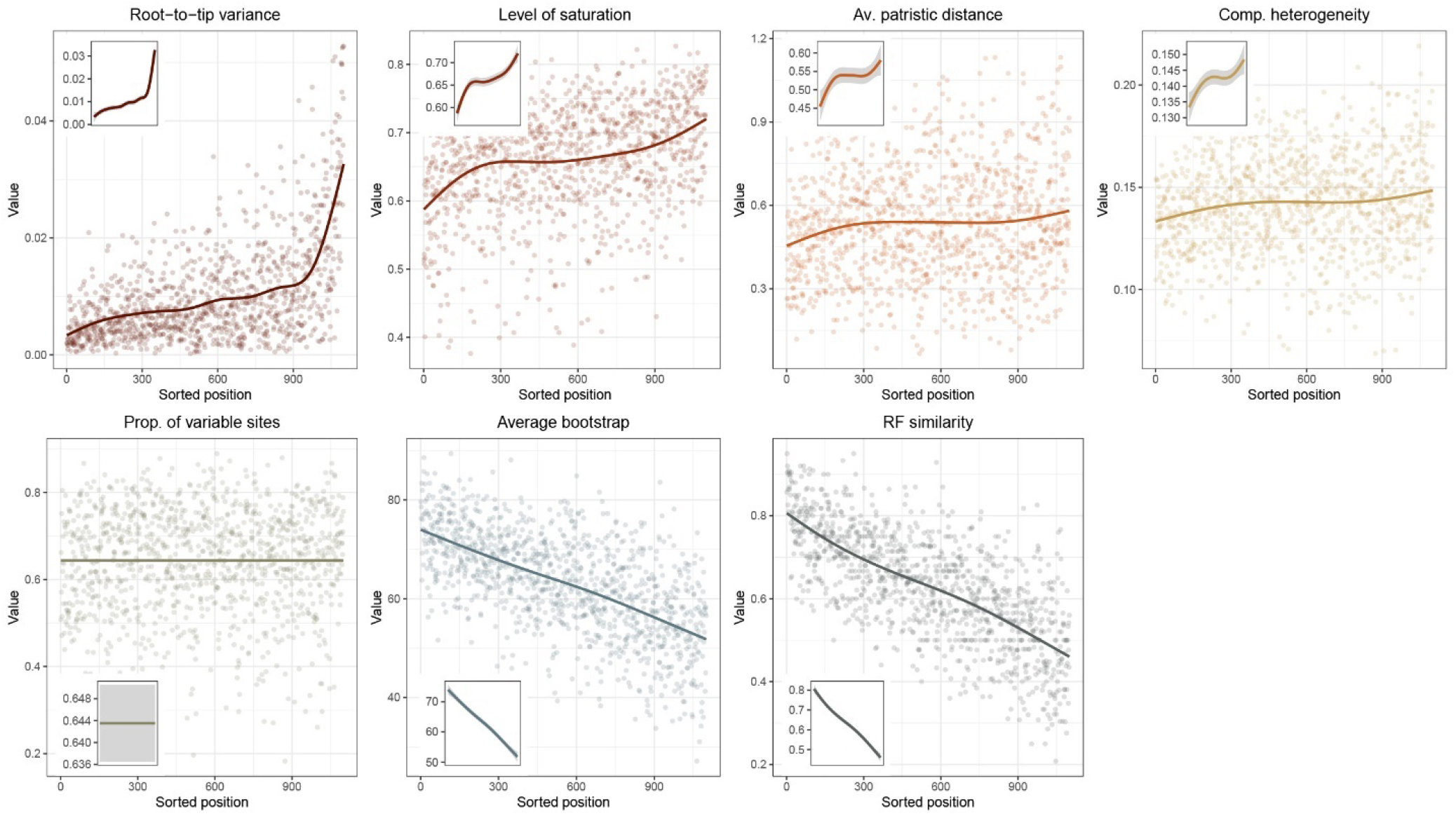
Distribution of gene properties after sorting the dataset for phylogenetic usefulness. Only ingroup taxa were considered for the estimation of these variables. Sorting was performed on PC 2, which explained 18.92% of total variance. Highly useful loci show limited evidence of suffering from systematic biases (top row), while having high phylogenetic signal (bottom row). The proportion of variable sites does not strongly load onto this axis.

**Figure S2:**
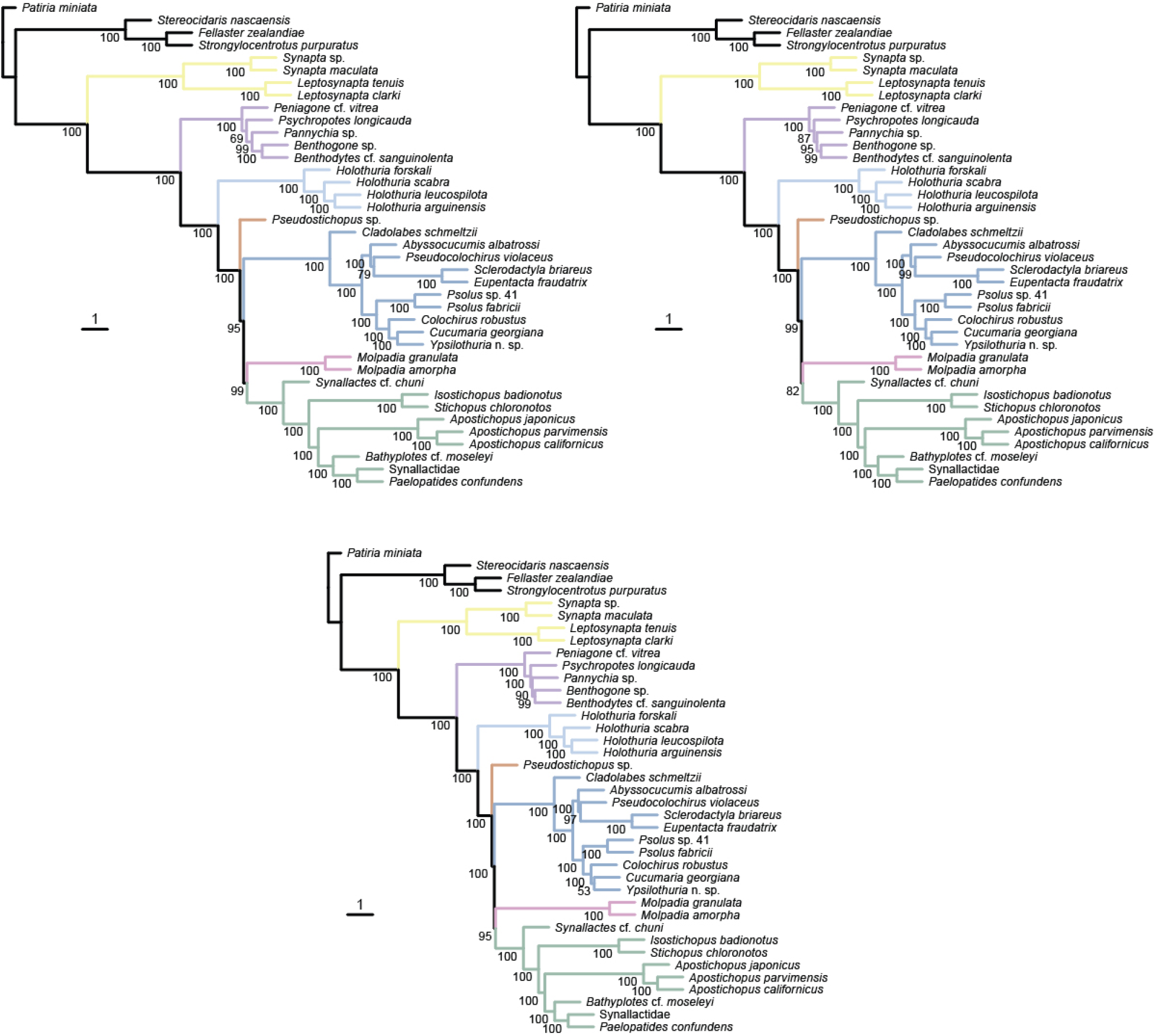
Topologies inferred using ASTRAL-III using the most phylogenetically useful quarter and half of loci (top left and top right, respectively), as well as the full dataset (bottom). Node values represent local posterior probabilities scaled to a maximum value of 100. Major holothuroid clades are colored as in Figure 2. Note that external branch lengths are meaningless in ASTRAL trees.

**Figure S3:**
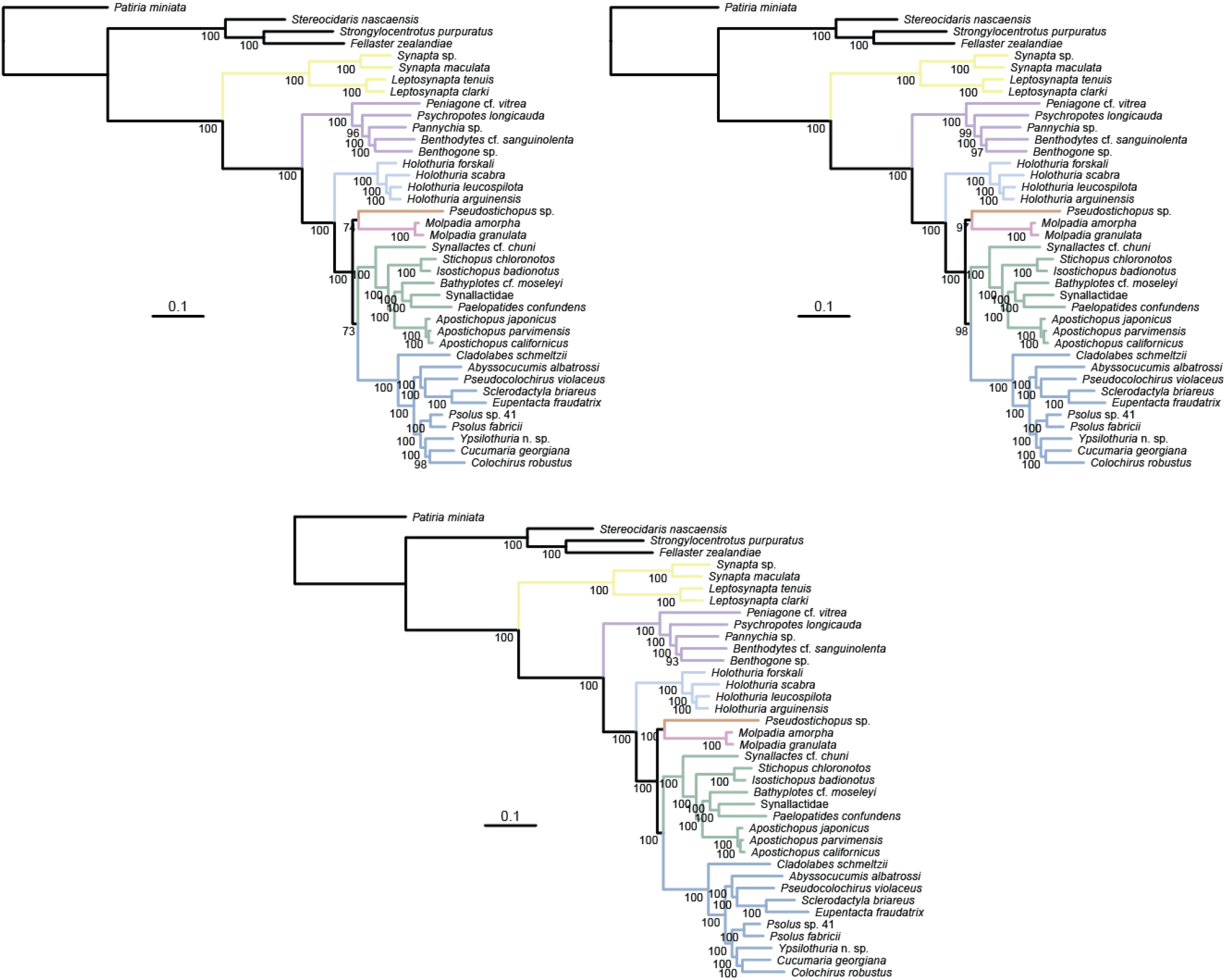
Topologies inferred using a best-fit partitioned model in IQ-TREE2 using the most phylogenetically useful quarter and half of loci (top left and top right, respectively), as well as the full dataset (bottom). Node values represent ultrafast bootstrap frequencies. Major holothuroid clades are colored as in Figure 2.

**Figure S4:**
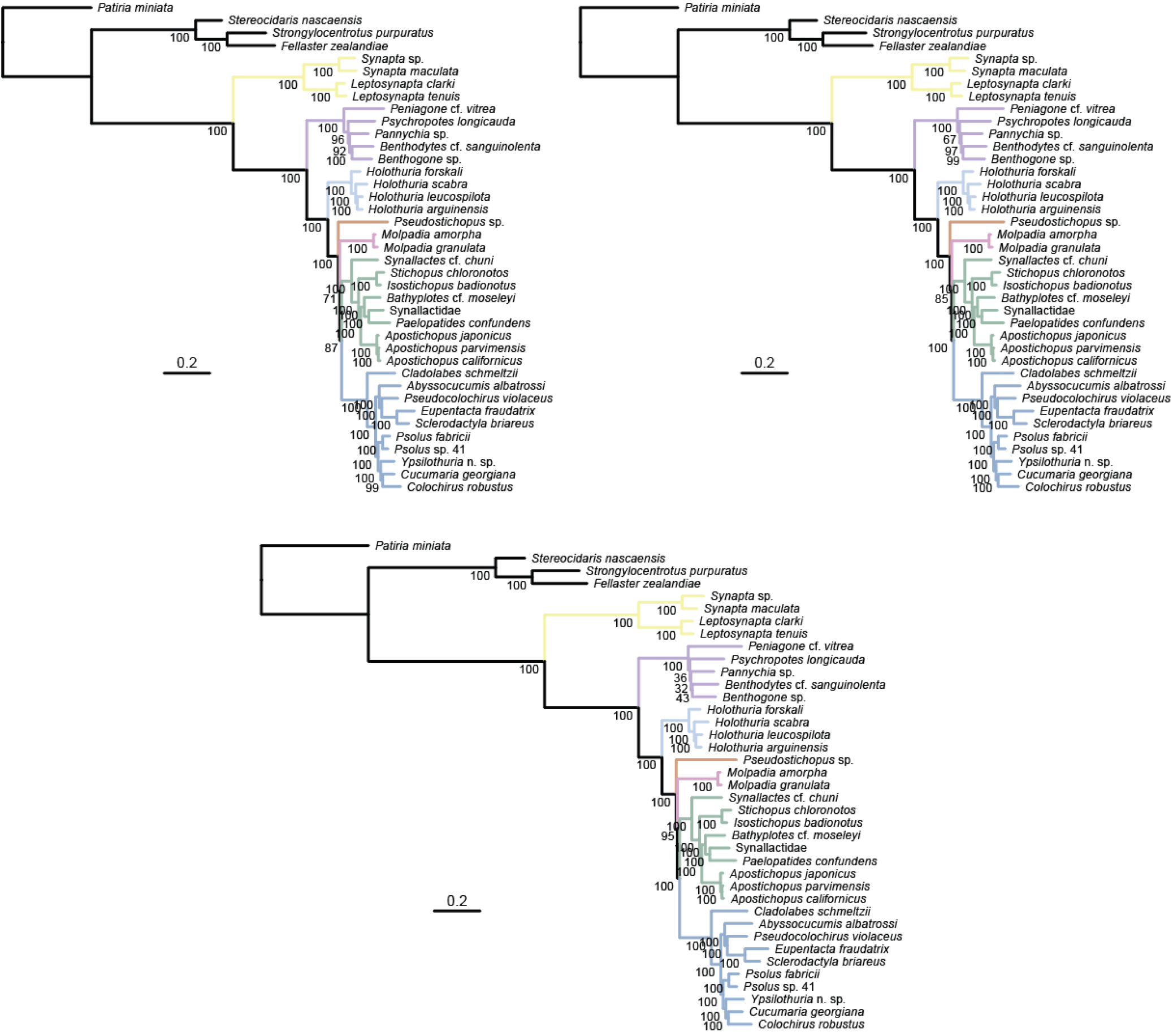
Topologies inferred using the site-heterogeneous model GTR+CAT-PMSF in IQ-TREE2 (with exchangeabilities and site-specific frequencies fixed to those obtained with PhyloBayes) using the most phylogenetically useful quarter and half of loci (top left and top right, respectively), as well as the full dataset (bottom). Node values represent ultrafast bootstrap frequencies. Major holothuroid clades are colored as in Figure 2.

**Figure S5:**
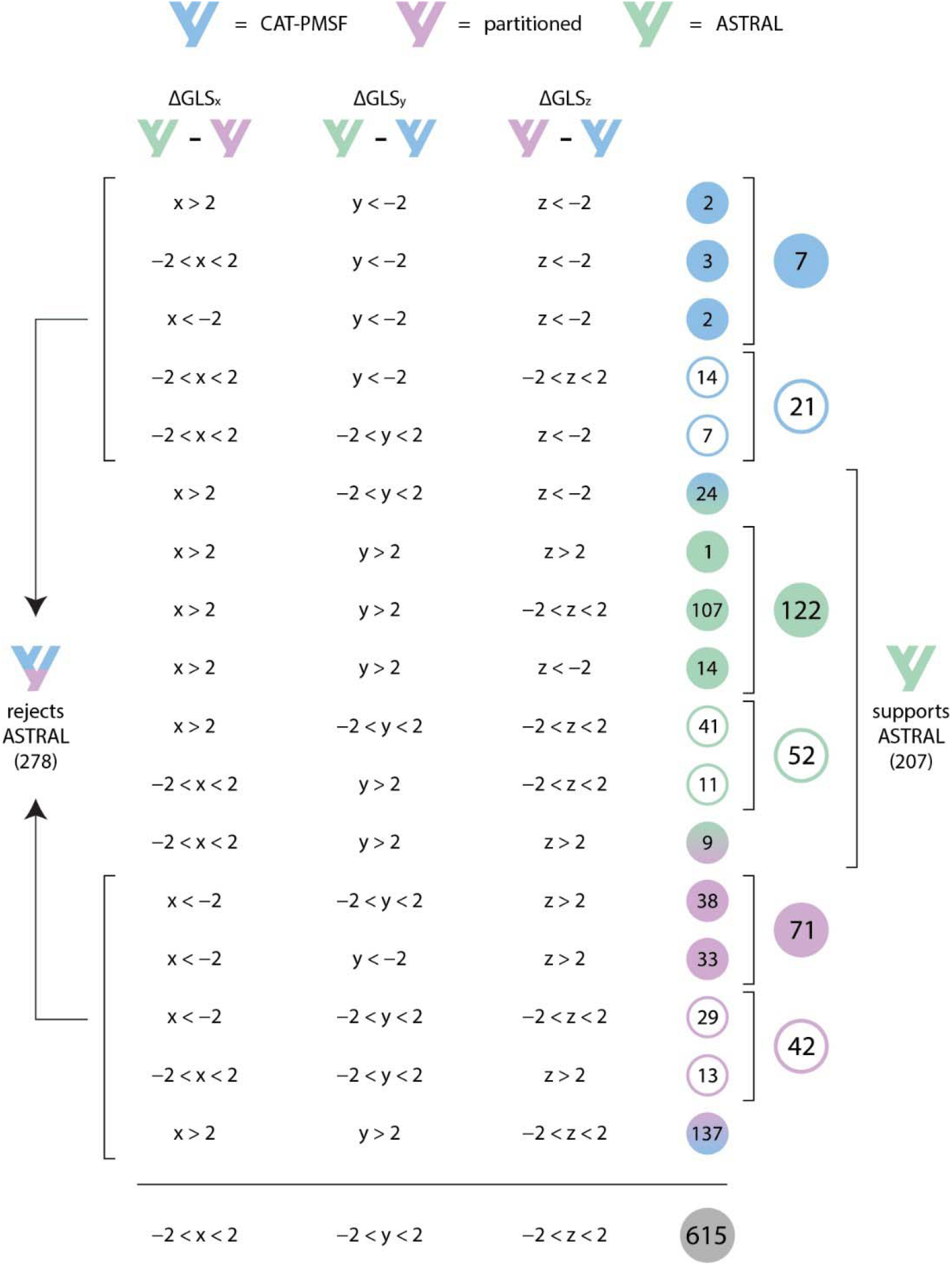
Categorization of loci following the application of ΔGLS thresholds of ± 2 log-likelihood units. Although 27 possible combinations exist, only 18 are realized. Filled and empty circles represent total or partial support for a given topology, respectively. Circles with a color gradient denote loci that provide support for two topological alternatives (i.e., consistently reject the third option). This same data is further summarized in Figure 4A.

**Figure S6:**
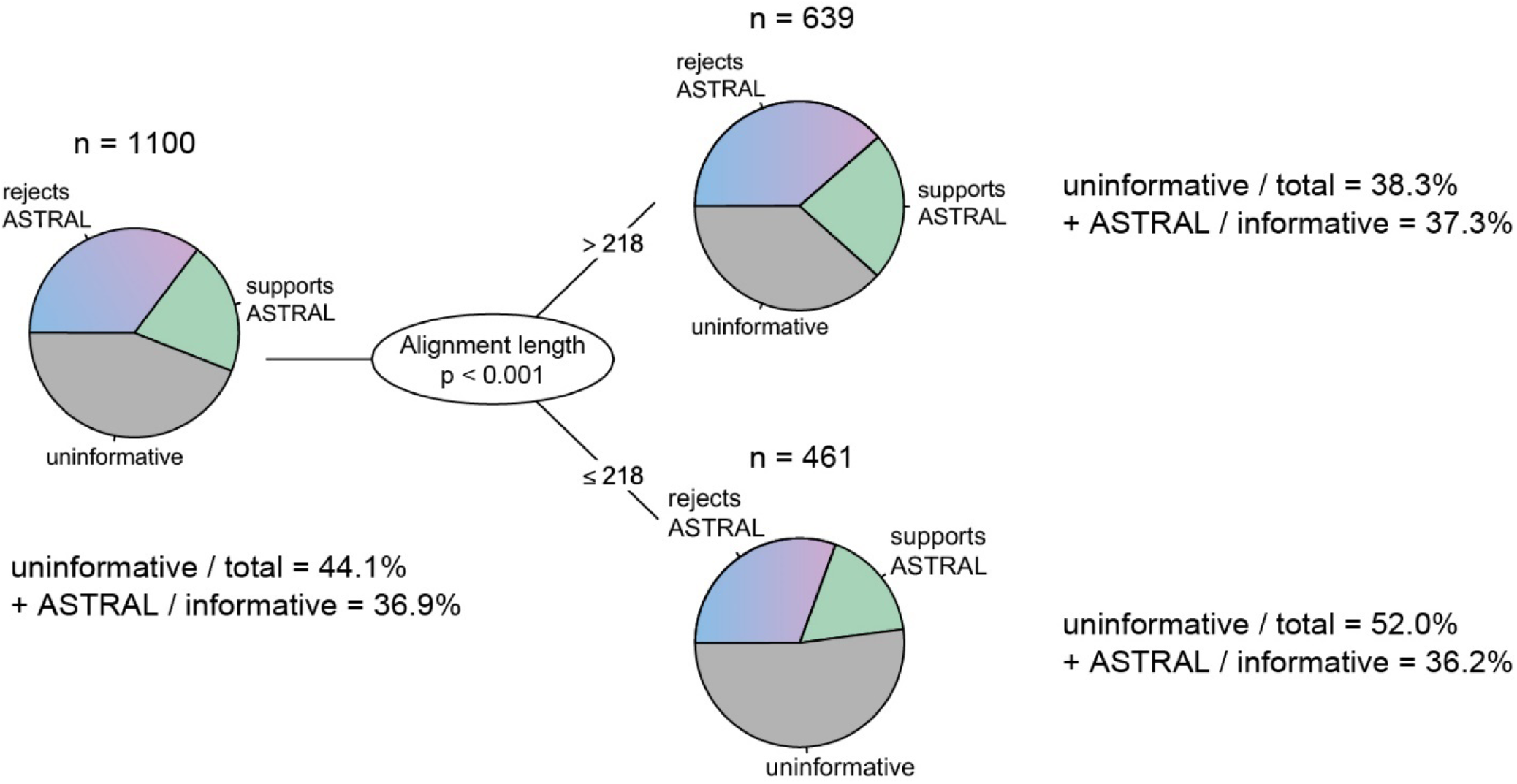
Results of the analysis of topological determinants using a classification tree. A single variable, alignment length, was found to be significantly associated with whether loci support or reject the ASTRAL resolution of Neoholothuriida, or remain topologically uninformative. Partitioning the data into small and large loci (with an automatically detected threshold of 218 base pairs) results in subsets that are enriched and depleted, respectively, in uninformative loci. The relative support for/against the topological alternatives, however, is not modified by this partitioning.

**Table 1:** Results of the exploration of topological determinants. Fifteen gene properties obtained with *genesortR* (see [38] for a definition of each) where correlated against the PC scores of ΔGLS. The magnitude of correlation against each is expressed as the endpoint of a vector of length = 1. R^2^ values represent squared correlation coefficients, *p*-values were obtained using 10,000 random permutations.

| Gene property | Type | Correlation<br>against PC 1 | Correlation<br>against PC 2 | $R^2$ | $p$-value |
| --- | --- | --- | --- | --- | --- |
| Root-to-tip variance | Bias | -0.012 | -1.000 | 0.002 | 0.311 |
| Level of saturation | Bias | -0.469 | -0.883 | 0.001 | 0.731 |
| Average pairwise patristic distance | Bias | -0.393 | -0.919 | 0.002 | 0.317 |
| Compositional heterogeneity (RCFV) | Bias | 0.123 | -0.992 | 0.001 | 0.489 |
| Average bootstrap support | Signal | -0.547 | -0.837 | 0.000 | 0.969 |
| Robinson-Foulds similarity | Signal | -0.122 | -0.993 | 0.000 | 0.908 |
| Phylogenetic usefulness<br>(PC 2 of *genesortR*) | Signal | -0.052 | -0.999 | 0.000 | 0.754 |
| Proportion of missing data | Information | -0.111 | 0.994 | 0.000 | 0.951 |
| Tree length | Information | -0.344 | -0.939 | 0.002 | 0.270 |
| Treeness | Information | 0.747 | 0.664 | 0.001 | 0.613 |
| Alignment length | Information | -0.147 | 0.989 | 0.001 | 0.689 |
| Level of occupancy | Information | 0.272 | 0.962 | 0.000 | 0.962 |
| Proportion of variable sites | Information | -0.490 | -0.872 | 0.001 | 0.460 |
| Tree length divided by number of<br>terminals | Rate | -0.397 | -0.918 | 0.002 | 0.278 |
| PC 1 of *genesortR* | Rate | -0.238 | -0.971 | 0.001 | 0.476 |

